# Bacterial metabarcoding reveals plant growth promoting members of the core *Brachypodium distachyon* root-associated microbiome overlooked by culture dependent techniques

**DOI:** 10.1101/2024.06.15.599150

**Authors:** Carl Otto Pille, Zahra F. Islam, Helen L. Hayden, Elena Colombi, Lok Hang Chan, Ute Roessner, Deli Chen, Hang-Wei Hu

## Abstract

Biofertilisers comprised of plant growth promoting bacteria (PGPB) present a promising sustainable alternative to synthetic fertilisers. Bacteria which consistently colonise roots of specific plants across distinct environments, known as that plant’s core root microbiome, are particularly promising due to their colonisation competency. However, traditional, culture-based techniques can overlook promising PGPB which do not display commonly screened for plant growth promoting traits. Although numerous studies have isolated beneficial root bacteria, few have combined bacterial metabarcoding with culture-based techniques to identify novel biofertiliser candidates. In a two-pronged approach, 16S rRNA amplicon sequencing was used to define the core root microbiome of the model cereal plant, *Brachypodium distachyon,* grown in four distinct soils. From 7,042 amplicon sequence variants (ASVs) detected in root fractions, only 40 ASVs were common at a prevalence of 80%. Core ASVs primarily belonged to the class *Alphaproteobacteria,* with the remainder comprising *Actinobacteria, Bacilli, Chloroflexia, Gammaproteobacteria* and *Negativicutes.* Secondly, *B. distachyon* root-associated bacterial strains were isolated from plants grown in the aforementioned soils. Of 207 root-associated isolates, 10 were identified as members of the core root microbiome, with the majority not displaying commonly screened for plant growth promoting traits. However, in a semi-hydroponic system, a core *Bacillus* and *Rhodococcus* strain significantly increased *B. distachyon* shoot dry weight by 32.8% and 40.0%, respectively. Additionally, two core *Bacillus* strains significantly increased root dry weight by 79.7 and 52.3%. This study demonstrates the potential of incorporating additional criteria afforded by culture-independent methods to screen for novel biofertiliser candidates typically overlooked by culture-dependent techniques.

**Importance:** Biofertilisers comprised of plant growth promoting bacteria (PGPB) present a sustainable, environmentally friendly alternative to synthetic fertilisers. The search for PGPB has historically relied on culture dependent methods to shortlist candidates by screening for plant growth promoting traits, such as N fixation or production of the plant hormone indole-3-acetic acid before conducting *in planta* trials. However, these screens may overlook promising bacterial strains which might promote plant growth via indeterminant mechanisms. Culture independent methods can take a snapshot of an entire microbiome without the need for culturing bacteria, allowing for additional selection criteria to be used when shortlisting PGPB. Our results show that targeting bacterial taxa which consistently colonise the roots of the model grass *Brachypodium distachyon* allowed for selection of bacterial isolates which were overlooked by culture-based PGPB trait screens. This result highlights the utility of combining traditional, culture-dependent methods with culture-independent techniques when searching for novel biofertiliser candidates.

## Introduction

As the global human population continues to grow, increased agricultural output from limited arable land has become vital (1). Sustainable alternatives to traditional synthetic fertilisers are desirable, as these agrochemicals often lead to detrimental downstream effects such as eutrophication of water bodies, release of greenhouse gases (2), and loss of soil biodiversity (3). Biofertilisers, made up of root-associated, plant growth promoting bacteria (PGPB), are considered a potential sustainable avenue for promoting crop growth (4). Biofertilisers contribute to improved plant growth through various mechanisms, such as improved nutrient delivery, phytohormone production, phytopathogen antagonism and abiotic stress resistance (5). Previous studies have shown that PGPB primarily belong to several genera, notably *Bacillus, Pseudomonas, Herbaspirillum, Azospirillum, Sphingomonas, Klebsiella, Stenotrophomonas* and *Burkholderia* (6). However, the translation of plant growth promotion by microbes observed in laboratory settings to the field remains challenging. A major cause of this phenomenon is fierce competition between inoculants and indigenous microbes for root colonisation (4). The root surface and its immediate surroundings, known as the rhizosphere, become battlegrounds for colonisation due to the presence of energy-rich compounds like sugars and amino acids in root exudates (7). Understanding and addressing these intricacies is crucial for optimizing the efficacy of biofertilizers in real-world agricultural scenarios.

Traditional culture-dependent techniques rely on the researcher’s discretion to initially isolate distinct bacterial strains, based on colony morphology and subsequent shortlisting of candidates following biochemical screens (8). However, focusing solely on strain performance in common biochemical screens may exclude strains that elicit their plant growth promoting effects through less common or unknown mechanisms. In a paradigm shift, recent studies have identified promising, novel biofertiliser candidates by first exploring the entire root microbiome via metabarcoding of the bacterial 16S rRNA gene (9–11). This method allows for a holistic view of the root microbiome, enabling the identification and selection of diverse taxa based on specific criteria, such as increased presence on and around roots under abiotic stress (12). One strategy in the formulation of biofertilisers involves incorporating bacteria that consistently colonise the roots of a particular plant across time and space, defining them as “core” members of that plant’s root microbiome. Core root-associated taxa are proficient competitive root colonisers due to their enduring presence across distinct environments in the presence of distinct indigenous microbiomes (13). Consequently, holistic identification of biofertiliser candidates requires the synergistic integration of molecular microbial ecology, traditional culture-dependent microbiology and plant phenotyping.

*Brachypodium distachyon* has been increasingly used as a model cereal in plant-microbiome interaction studies due to its small size and relatively analogous responses to biotic and abiotic stimuli as important crops such as barley and wheat (14, 15). As studies have reported that domesticated cereal crops may have lost some ability to communicate with and attract beneficial soil bacteria, *B. distachyon* may also be considered a good PGPB reservoir due to its status as a wild plant (16, 17). Several studies have investigated the root-associated microbiome of *B. distachyon* (18, 19), though none have defined its core root-associated microbiome nor isolated its root-associated bacteria. The environment plays a greater role in shaping the root-associated microbiome compared to factors such as plant genotype or development stage (20–22). Thus, this study aimed to explore the *B. distachyon* root associated microbiome across four soils chosen for their distinctly different land uses.

Specifically, the objectives of this study were: (1) to define the core *B. distachyon* root associated microbiome via 16S rRNA gene metabarcoding; (2) isolate core root-associated taxa and characterise their plant growth promoting potential through common biochemical screens; and (3) determine the plant growth promoting efficacy of core isolates on *B. distachyon* biomass in a semi-hydroponic, gnotobiotic system.

## Results

### Microbiome composition

*B. distachyon* plants were grown in four distinct soils. The first two soils were collected in urban grassland in Royal Park (RP) and remnant scrubland along a Train line (TL) in Melbourne, Victoria. The remaining two soils were collected from managed cropland (DA) and unmanaged grassland (DR) at The University of Melbourne agricultural campus in Dookie, Victoria. Soils varied in their nitrate (2.9 – 52 mg/kg), ammonium (1.7 – 57 mg/kg), K (12 – 340 mg/kg) and P (7.7 – 130 mg/kg) contents, as well as pH (4.2 – 5.2) and organic matter (2 – 12%, Table A1). Samples were separated into bulk soil, rhizosphere soil and root-associated fractions before investigating their microbial communities (for each fraction of each soil n = 6 except RP root-associated fraction where n = 5).

After filtering out low quality and non-prokaryotic reads, 25,290 amplicon sequence variants (ASVs) were classified across all fractions, with 7,042 ASVs observed in root-associated fractions. At the phylum level, bulk soil fractions were dominated by *Actinobacteriota* (55.4 – 64.4%), *Proteobacteria* (13.3 – 16.5 %), *Chloroflexi* (2.6 – 13.1%), *Acidobacteriota* (5.4 – 10.19 %) and *Firmicutes* (0.6 – 13.14%) (Fig. 1a). At the order level, bulk soil fractions were mainly composed of *Solirubrobacterales* (11.4 – 20.4%), *Frankiales* (5.8 – 26.9%), *Rhizobiales* (3.9% - 9.6%), *Gaiellales* (3.2 – 17.0%) and *Bacillales* (0.3 – 12.3%). DA bulk soils harboured a much larger proportion of *Bacillales* (12.3%) than other bulk soils (0.3 - 2.09%). Additionally, TL bulk soils contained a larger proportion of *Frankiales* (26.9%) and a lower proportion of *Gaiellales* (3.2%) than other bulk soils (5.8 – 15.4% & 8.4 - 17.9%, respectively) (Fig. 1b).

**Figure 1:**
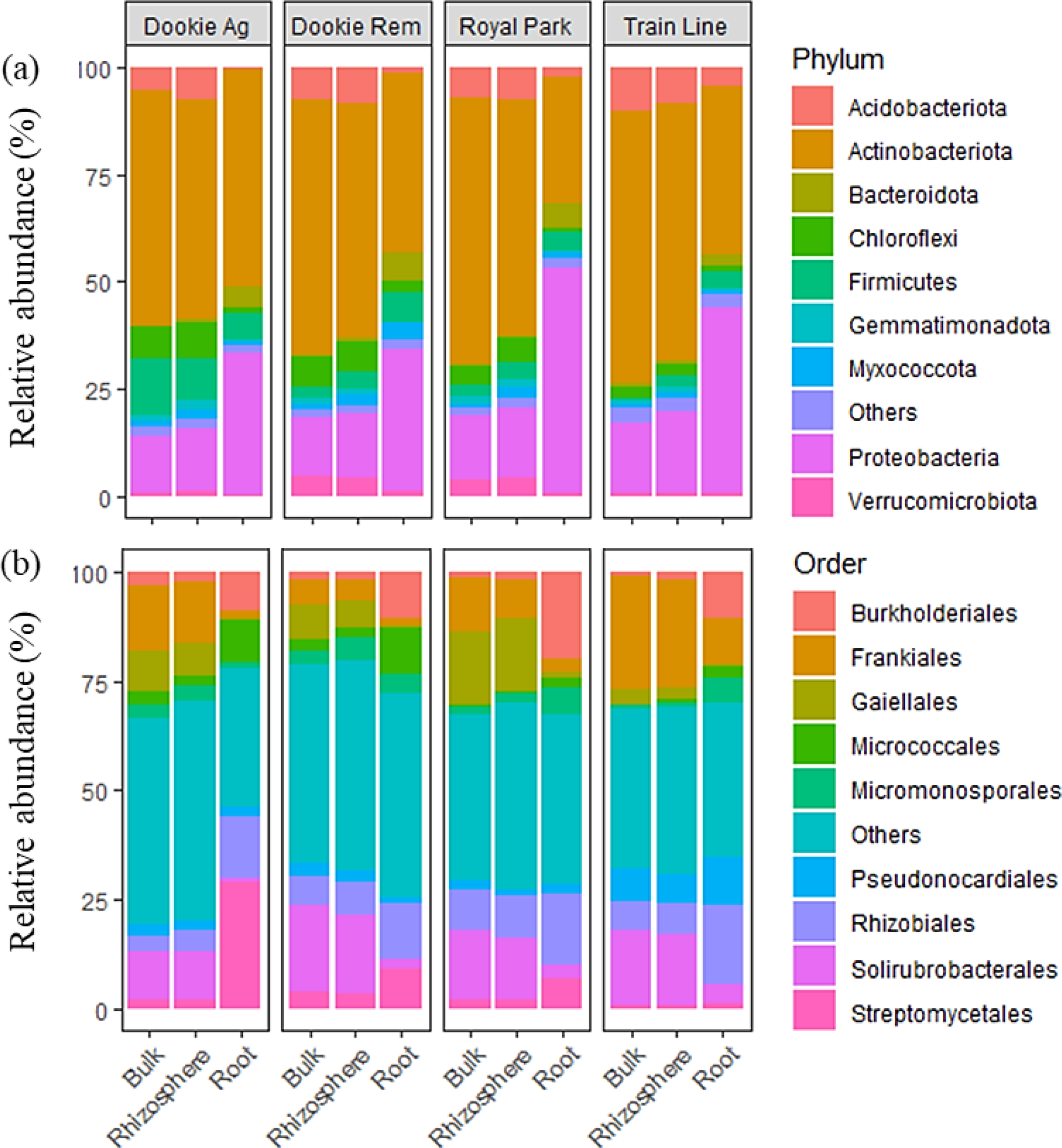
Relative abundance of bacterial taxa in bulk soil, rhizosphere soil and root fractions of *B. distachyon* grown in managed Dookie soil (Doogie Ag), unmanaged Dookie soil (Dookie Rem), Royal Park soil and Train Line soil. Composition presented at phylum (a) and order (b) levels.

In comparison to bulk soil fractions, *Actinobacteriota* were less pronounced (30.1 – 49.8 %) in root-associated fractions, while *Proteobacteria* were much more relatively abundant (33.7 – 52.7%). *Firmicutes* (4.3 – 6.5%) and *Bacteroidota* (2.8 – 6.6%) made up the majority of the remaining phyla in root-associated fractions (Fig. 1a). At the order level, root-associated fractions were made up of *Rhizobiales* (13.0 – 18.6%)*, Streptomycetales* (1.3 – 28.43%) and *Burkholderiales* (9.4 – 19.9%, Fig. 1b). DA root-associated fractions harboured a much larger proportion of *Streptomycetales* (28.43%) than other soils (1.3 – 9.17%). Additionally, *Streptomycetales* were more relatively abundant in DA root-associated fractions than bulk soil fractions, while this trend was reversed in all other soils (Fig. 1c). Finally, rhizosphere soils were very similar to bulk soil fractions at both the phylum and order level in all soils (Fig. 1a and b).

### Microbiome diversity

Alpha diversity analysis based on the Shannon diversity metric showed that root fractions were significantly less diverse than bulk and rhizosphere soil fractions, regardless of soil type (p < 0.05, Fig. 2a). The largest reduction in diversity between bulk soil and root-associated fractions was observed in DA samples.

**Figure 2:**
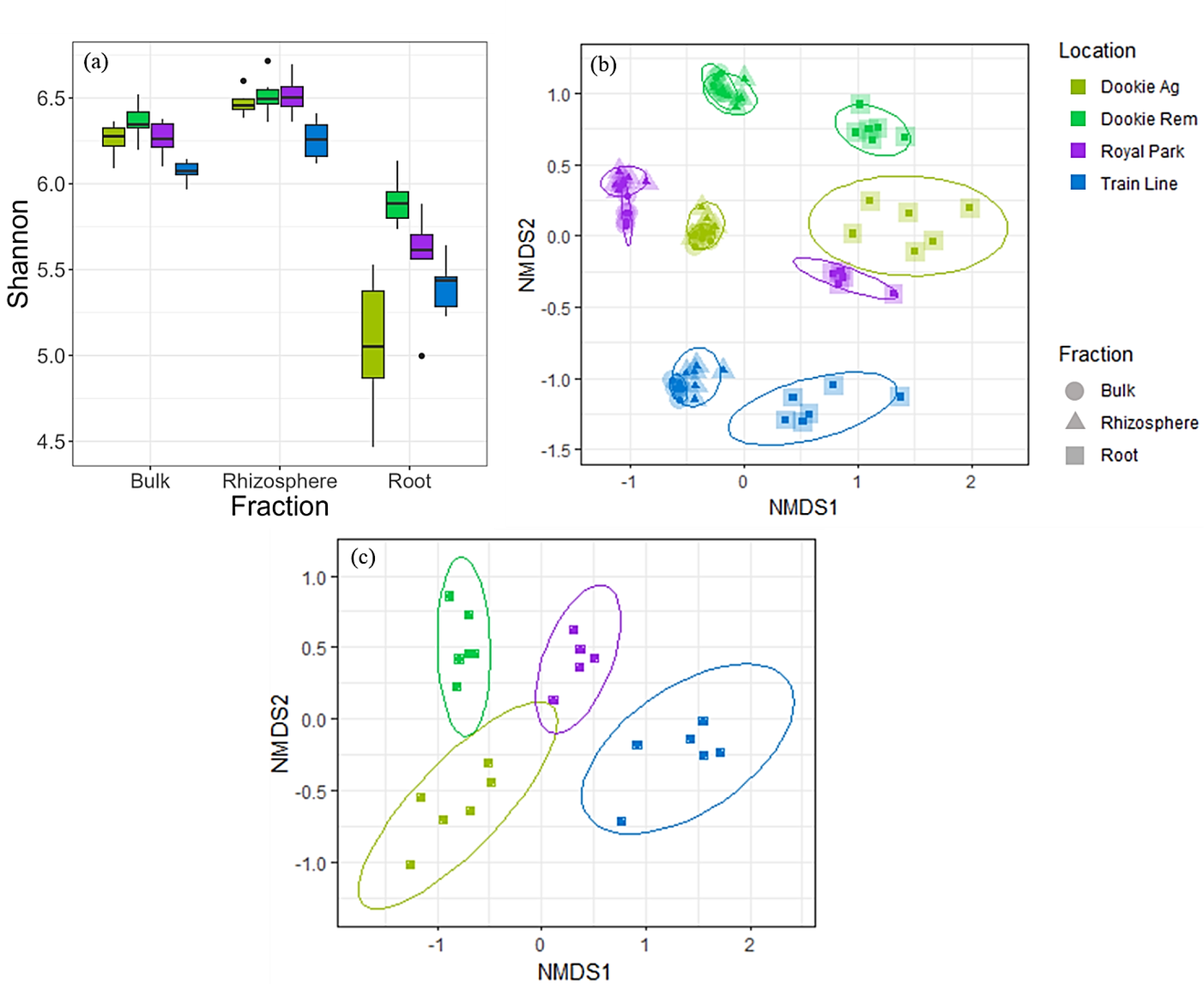
(a) Shannon diversity of bulk soil, rhizosphere soil and root microbiome fractions of *B. distachyon* plants grown in four contrasting soils. (b) Non-metric multidimensional scaling (nMDS) ordination of the Bray-Curtis dissimilarity matrix of all samples with colours and shapes representing soil type and fraction, respectively (Bray-Curtis stress = 0.183). (c) Non-metric multidimensional scaling (nMDS) ordination of the Bray-Curtis dissimilarity matrix of root samples (Bray-Curtis stress = 0.106).

To find differences in bacterial communities between samples, beta diversity analysis was conducted to find differences between samples based on Bray-Curtis dissimilarity and visualised with an NMDS ordination (Fig. 2b and c). Bulk and rhizosphere soil fractions clustered together and away from root fractions (Fig. 2b). When only comparing root-associated fractions, samples clearly clustered based on location (Fig. 2c).

Permutational multivariate analysis of variance (PERMANOVA) analysis showed that location was a larger determinant of microbiome composition than fraction type. This effect was most pronounced in bulk soil and rhizosphere soil fractions, having a lesser effect in root-associated fractions (Table A2). Pairwise comparisons showed that bulk soils from each location were significantly different from each other, as were root-associated fractions (p < 0.001).

### The core *Brachypodium distachyon* root-associated microbiome

Compared with similar studies (21, 23), this work selected a conservative prevalence threshold of 80% as richness remained stable at higher threshold values (Fig. A1). Of 7,042 ASVs observed in root samples, 40 ASVs were common in 80% of root samples. In root-associated fractions, the sum of the relative abundances of core root-associated ASVs in DA, DR, RP and TL soils was 16.4, 16.1, 18.0 and 18.4%, respectively. *Xanthobacteracea* ASV_2, *Streptomyces* ASV_7 and *Rhodococcus* ASV_72 were the most relatively abundant core ASVs in root associated fractions (Fig. 3b).

**Figure 3:**
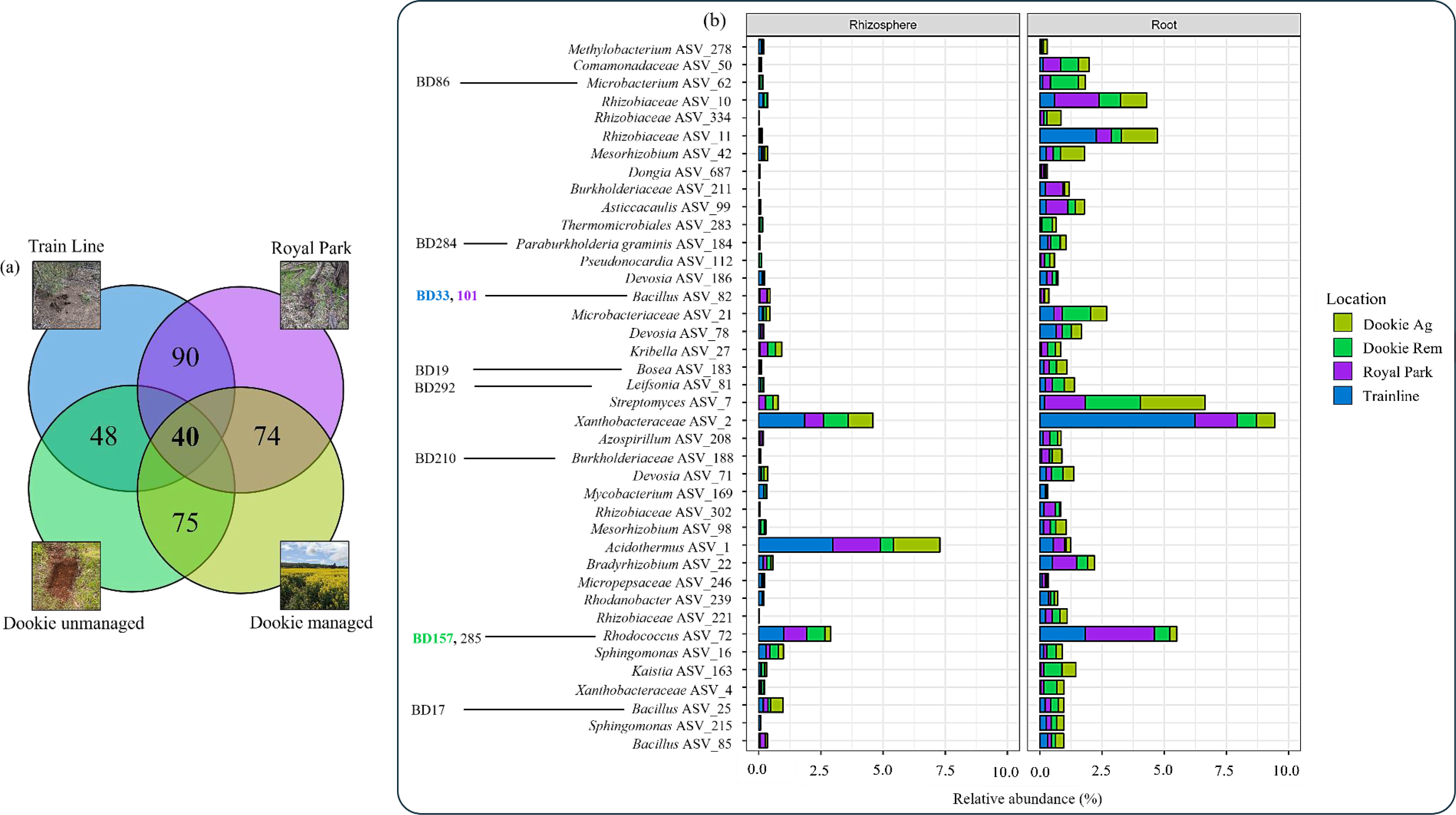
(a) Number of ASVs common between root fractions of *B. distachyon* grown in four soils at a prevalence of 80%. (b) Relative abundance of bacterial classes making up the core *B. distachyon* root microbiome. Isolates with “BD#” identifiers are listed beside their respective core ASVs, with those in bold observed to promote *B. distachyon* shoot and/or root growth.

The majority of core root-associated ASVs were classified as belonging to the *Proteobacteria, Actinobacteriota* or *Firmicutes* phyla. Notable genera previously reported as PGPB included: *Bacillus, Bradyrhizobium, Sphingomonas, Burkholderia* and *Azospirillum*. The relative abundance of individual core root ASVs varied between locations, with most core ASVs comprising less than 1% of the root-associated microbiome of *B. distachyon* in each soil. Apart from *Acidothermus* ASV_1, core ASVs were more abundant in root fractions than rhizosphere fractions (Fig. 3b).

### Isolation and characterisation of *B. distachyon* root associated bacteria

A total of 207 bacteria were isolated from the roots of *B. distachyon* grown in the four soils and were identified via Sanger sequencing of 16S rRNA amplicons from pure cultures. Overall, 37 genera were isolated, with *Pseudomonas* and *Microbacterium* species most commonly isolated (Fig. A2).

The 16S rRNA sequences from isolates were aligned with core ASV sequences to determine whether any isolates displayed 100% identity with core ASV sequences and could thus be considered core isolates. Core isolates with identical sequences but isolated from different soils were considered unique core isolates. Ten of the 207 isolates were identified as unique core isolates, representing eight core ASVs with at least one core isolate originating from each soil. These core isolates belonged to the genera *Neobacillus, Bacillus, Bosea, Burkholderia, Paraburkholderia, Leifsonia, Microbacterium* and *Rhodococcus*. Biochemical screens showed that *Neobacillus* BD17 and *Bosea* BD19 could produce indole compounds, while *Burkholderia* BD284 could solubilise Ca_3_(PO_4_)_2_ (Table 1). None of the core isolates were observed to fix N_2_ or produce siderophores (Table 1).

**Table 1:** Biochemical screens of core *B. distachyon* root associated isolates.

| Isolate (BD#) | Origin* | N<sub>2</sub><br>fixation | Siderophore<br>production | Phosphate<br>solubilisation** | IAA<br>production<br>(µg/mL) |
| --- | --- | --- | --- | --- | --- |
| *Bacillus* BD17 | TL | - | - | - | 17 |
| *Bacillus* BD33 | TL | - | - | - | - |
| *Bacillus* BD101 | RP | - | - | - | - |
| *Bosea* BD19 | TL | - | - | - | 26 |
| *Paraburkholderia*<br>BD210 | DU | - | - | - | - |
| *Burkholderia*<br>BD284 | DM | - | - | ++ | - |
| *Leifsonia* BD292 | DM | - | - | - | - |
| *Microbacterium*<br>BD86 | RP | - | - | - | - |
| *Rhodococcus*<br>BD157 | DU | - | - | - | - |
| *Rhodococcus*<br>BD285 | DM | - | - | - | - |

IAA = Indole acetic acid
\*RP = Royal Park, TL = Train line, DM = Dookie managed, DU = Dookie unmanaged
\*\* Phosphate solubilisation index (PSI) = Zone of clearing diameter (D<sub>h</sub>)/Colony diameter (D<sub>c</sub>)
PSI = 1 (+), 2 (++), 3 (+++)

### *B. distachyon* growth promotion by core isolates

The ten unique, core isolates previously screened for common plant growth promoting traits were then tested for *B. distachyon* growth promoting ability in a gnotobiotic, semi-hydroponic system. *Bacillus* BD33 and *Rhodococcus* BD157 significantly increased shoot dry weight by 32% and 45%, respectively (p < 0.05, Fig. 4). *Bacillus* BD33 and *Bacillus* BD101 also significantly increased root dry weight by 79% and 52%, respectively (p < 0.05, Fig. 4). Additionally, *Bacillus* BD33 significantly increased leaf number by 28% (p < 0.05), and *Bosea* BD19 significantly altered biomass allocation towards roots, rather than shoots (p < 0.05, Table A3).

**Figure 4:**
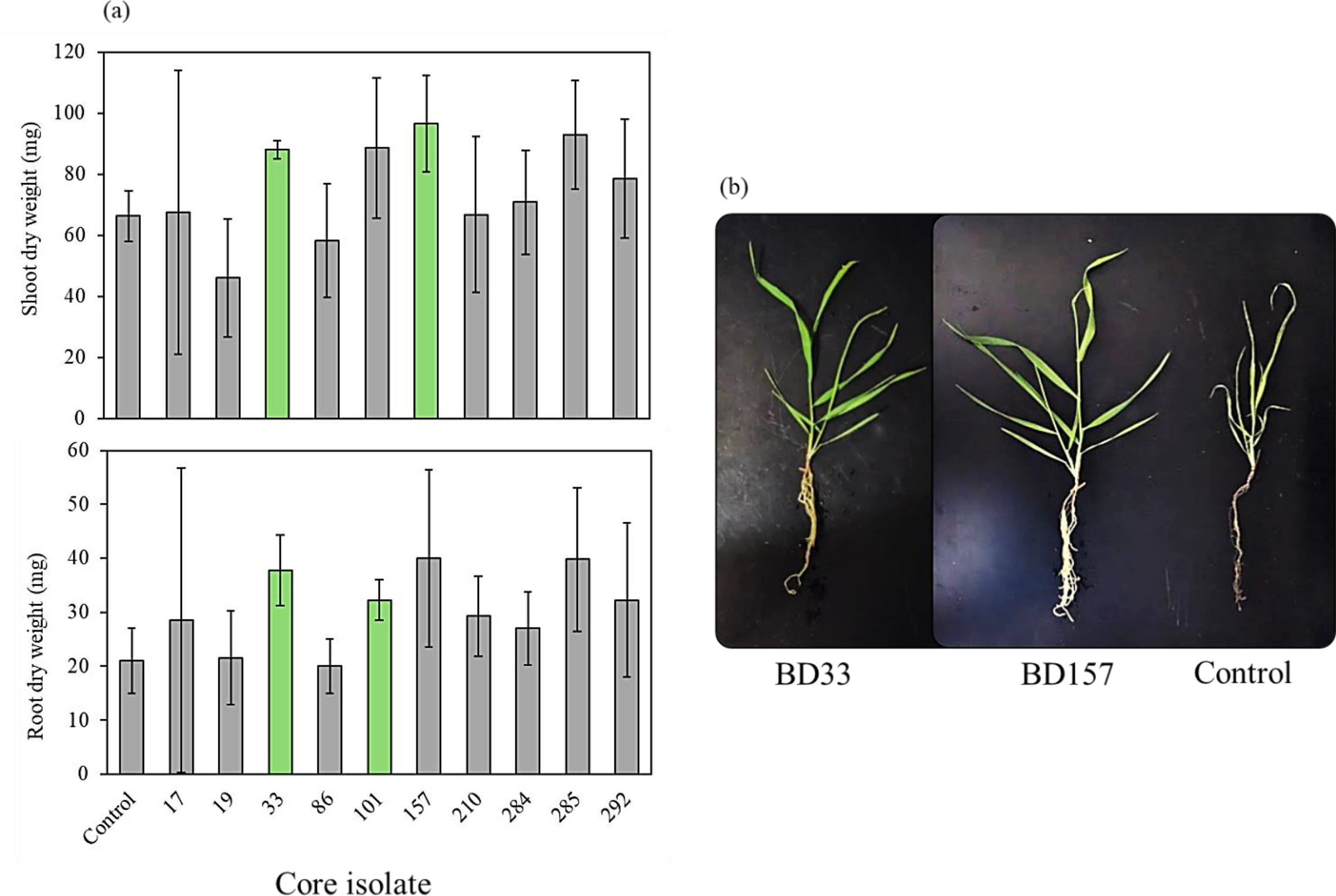
(a) Mean root and shoot dry weights of *B. distachyon* plants inoculated with core isolates and associated 95% confidence intervals (* = p <0.05, n = 4). (b) *Brachypodium distachyon* inoculated with *Bacillus* BD33, *Rhodococcus* BD157 and control.

## Discussion

This study identified the core root microbiome of *B. distachyon* grown in four distinct soils. Of 207 root associated bacterial isolates, 10 were identified as unique core isolates, with a core *Rhodococcus* and two core *Bacillus* isolates observed to significantly increase *B. distachyon* shoot or root dry weight (Fig. 3 and 4).

The four soils used in this study differed in their land use histories and physiochemical properties (Table 1), which have previously been identified as significant drivers of bacterial diversity (24) and stronger determinants than plant developmental stage or genotype in shaping plant microbiome diversity (25). Subsequently, bacterial communities among bulk soil fractions were found to be distinct in each sampled soil, indicating their suitability for defining core *B. distachyon* root associated taxa persistent across contrasting indigenous microbiomes (Fig. 2). Root-associated fractions of *B. distachyon* grown in each soil had significantly lower bacterial diversity than bulk and rhizosphere soil fractions, consistent with previous studies (23). This was likely due to selective pressure exerted by root exudates (20). At the phylum level, the core *B. distachyon* root associated microbiome resembles those of agriculturally relevant grasses, such as wheat (18, 26, 27), composed primarily of *Actinobacteriota, Proteobacteria* and *Firmicutes*. Along with previous reports, this further supports the suitability of *B. distachyon* as a model for investigating plant-microbe interactions and isolation of potential biofertiliser candidates. Differences at lower taxonomic levels are likely due to soil-specific microbiome variations.

At the conservative prevalence threshold of 80% (21, 23), the core root-associated *B. distachyon* microbiome comprised 40 ASVs. Core ASVs accounted for 16 -18% of the relative bacterial abundance in root fractions (Fig. 3), indicating their dominance in this niche. Many of these core ASVs belong to genera previously reported for their plant beneficial effects, through indirect or direct growth promotion mechanisms. These include important nutrient cycling genera, such as the N_2_ fixing *Bradyrhizobium, Devosia* and *Sphingomonas*, as well as plant growth promoting and phytopathogen antagonists *Bacillus* and *Paraburkholderia* (28, 29). Eight core ASVs were represented by 10 isolates, with two unique core isolates from different soils found to match with *Bacillus* ASV_82 and *Rhodococcus* ASV_72. However, the majority of core ASVs were not isolated, likely due to suboptimal growth conditions for these specific taxa. In the future, rather than con-currently isolating bacteria with general, nutrient rich growth media, culture-independent analysis should be followed by targeted isolation of taxa of interest with more appropriate media and culture conditions (11).

Almost none of 10 core isolates displayed commonly screened for plant growth promoting traits (Table 1). While it is possible that core ASVs which did not display identity with isolates possess the screened traits, they may also indirectly affect plant growth or play roles in microbiome assembly, structure and function. They may also directly antagonize phytopathogens or increase plant resistance to abiotic and biotic stressors (30), or only express these traits under specific environmental conditions. Moreover, genera of the core *B. distachyon* root associated microbiome identified in the current study, such as *Azospirillum*, *Bacillus, Bradyrhizobium* and *Burkholderia*, have been reported in several studies as components of core root associated microbiomes across multiple plant lineages such as ferns, lycopods, gymnosperms and angiosperms, suggesting the existence of a “universal” core root microbiome (20–22). *Bacillus* strains belonging to this “universal” core root microbiome have been isolated from tomato (*Solanum lycopersicum*) and found to promote wheat growth, suggesting that a core root microbiome has co-evolved with land plants and may be beneficial across relatively unrelated plant lineages (31).

The core isolates *Bacillus* BD33 and *Rhodococcus* BD157 significantly increased *B. distachyon* shoot dry weight, while *Bacillus* BD33 and *Bacillus* BD101 significantly increased root dry weight (Fig. 4). The genus *Bacillus* is frequently isolated from plant roots and observed to promote plant growth, including in *B*. *distachyon* (32, 33). *Bacillus* spp. are promising candidates for biofertilisers due to their high metabolic diversity, with single strains being shown to promote growth across various plant hosts and soil types (28, 33). Additionally, unlike other common plant growth promoting bacteria such as *Pseudomonas*, *Bacillus* are spore forming and can be stored for extended periods of time as spores. Due to these attributes, several commercially available biofertiliser products are made up of *Bacillus* strains (33).

In contrast, the genus *Rhodococcus* has been reported in close association with plant roots, though usually as the causative agents of diseases such as leafy gall disease in a range of plants and bushy top syndrome in pistachio (34, 35). However, some studies have found that *Rhodococcus* species can promote the growth of *Arabidopsis thaliana* and *Pisum sativum* in hydrocarbon and heavy metal contaminated soils (36, 37). To the best of our knowledge, this study is the first report of *Rhodococcus* promoting the growth of a monocot plant.

Interestingly, the mechanisms underlying the ability of *Bacillus* BD33 and BD101 and *Rhodococcus* BD157 to promote plant growth remain unclear, as these isolates did not possess any plant growth promoting traits that were screened for in this study, such as production of the auxin indole-3-acetic acid (Table 1). These isolates may produce other plant hormone homeostasis altering compounds, such as gibberellins or cytokinins (38). This result shows that reliance on biochemical screens may overlook promising novel biofertiliser candidates, which may be identified using criteria afforded only by culture-independent methods such as bacterial metabarcoding.

### Conclusions

This study has defined the core microbiome associated with *Brachypodium distachyon* roots and isolated several plant growth promoting bacteria belonging to its core root-associated microbiome. Due to their observed persistent root colonisation in the presence of distinct soil microbiomes, these core plant growth promoting isolates harbour great potential as biofertilisers. Further work will elucidate the mechanism underpinning core isolate plant growth promotion, their colonisation lifestyle, host range and ability to compete in the rhizosphere in the presence of a natural, complex microbiome. Finally, this study has highlighted that investigation of the entire microbiome is useful in identifying biofertiliser candidates which may go unnoticed using traditional, culture-based screening techniques.

### Methods and materials

### Plant materials and soil collection

Topsoil samples were collected from a depth of 0 – 30 cm at four distinct sites in September 2022 in Victoria, Australia. The first site, Royal Park in Melbourne (-37.790, 144.951; Royal Park (RP)), Australia is a managed park land subject to irregular treatments with pesticides and fertilisers. The second site is a strip of land adjacent to a train line in Parkville (-37.783, 144.944; Train line (TL)), Melbourne with a long history of native flora conservation. The third and fourth sites are a managed canola field (-36.377, 145.706; Dookie managed, DA) and unmanaged grassland (-36.382, 145.703; Dookie unmanaged, DR), respectively, at The University of Melbourne’s agricultural campus in Dookie, Victoria, Australia. The sites were selected for their varying land uses, anticipating the presence of distinct bacterial communities suitable for the identification of core bacteria. Soil samples from each site were placed in plastic bags and kept at 4℃ for short-term preservation.

### Soil physiochemical characterisation

Soils were sent to Nutrient Advantage, Werribee, Australia for physiochemical analysis according to National Association of Testing Authorities accredited standard procedures ^(^http://www.nata.com.au). Nitrate (NO_3-_) and ammonium (NH_4+_) were analysed using flow injection analysis (FIA), potassium (K) using inductively coupled plasma optical emission spectroscopy (ICP-OES), soil organic carbon content using UV spectroscopy, and soil pH in the presence of CaCl_2_ and phosphorus (P) using the BSES method.

### Soil and root sampling for bacterial community analysis

All plant work used the BD21-3 line of *Brachypodium distachyon. B. distachyon* seeds had their lemma removed before surface sterilisation with 70% ethanol and 5% sodium hypochlorite for 30 s and 5 min, respectively, before washing five times with sterile dH_2_O. Surface sterilised seeds were imbibed for 48 h at 4℃ in sterile dH_2_O. Imbibed seeds were germinated on 0.9% agar plates kept at room temperature in the dark for 72 h. Three germinated seeds were sown in a triangular arrangement in 14 cm × 12 cm (width × height) pots, with 6 pots per soil for a total of 24 pots. The plants were grown for 28 days in a Conviron growth cabinet under a 18/6 light cycle at 24℃ and 18℃, respectively. Plants were watered every 3 days with sterile MilliQ water.

Three plants per pot were harvested and pooled prior to fractionation and DNA extraction. Bulk soil was collected from the centre of each pot and stored at -80℃. Roots were harvested by squeezing the sides of the pot and carefully sliding out the root system. Very loosely attached soil was gently shaken off before placing the remaining roots and tightly adhered soil in a 50 mL Falcon tube containing 30 mL of 0.2 mM CaCl_2_. Falcon tubes were vortexed three times for 30 s, after which roots were removed and placed in 50 mL Falcon tubes containing sterile 1 × phosphate buffered saline (PBS). The remaining soil was pelleted by centrifuging at 5000 × g for 5 minutes and stored at -80℃, constituting the rhizosphere soil fraction. Falcon tubes containing roots were placed in an Elma S 300H sonication bath and sonicated for 5 min at 37 kHz. Subsequently, roots were removed and stored at -80℃, constituting the root-associated fraction.

### DNA extraction

Six root samples per soil (except RP where n = 5) were homogenised by submerging in liquid nitrogen and grinding with a sterile mortar and pestle, prior to DNA extraction. DNA was extracted from root and soil samples using a DNeasy PowerSoil Pro DNA isolation kit (QIAGEN) and a QIACUBE automated extraction system, according to the manufacturer’s instructions. The DNA extracts were screened for purity using a Nanodrop spectrophotometer before measuring DNA concentration using a Qubit fluorometer. DNA extracts (10 ng/µL) were sent to the Australian Genome Research Facility for amplicon sequencing of the 16S rRNA gene using the primers 341F (5’-CCTACGGGNGGCWGCAG ′) and 806R (5 ′ - GGACTACHVGGGTWT CTAAT′) (39).

### Bioinformatic analysis of 16S rRNA gene sequences

Raw sequence reads in the FASTQ format were pre-processed on the Spartan high-performance computer (HPC) at The University of Melbourne. Sequences with average quality scores < 20 were filtered out using DADA2 v1.26.0 (40). Adapters were trimmed before demultiplexing, denoising and mapping reads to the SILVA database v.138.1 (41) using QIIME2 v.2021.8 (42). Amplicon sequences with 97% similarity were assigned to the same amplicon sequence variant (ASV). The resulting high-quality sequences were used for downstream analysis in R v4.4.0 and R Studio v2023.06.0 (43). ASVs that did not align with any phylum or were classified as chloroplast, mitochondrial or eukaryotic were excluded from the dataset. ASVs with less than five total counts in each sample were also removed.

Sample sizes were normalised by rarefying each sample to 90% of the total count sum of the smallest sample. Shannon diversity was estimated using the “phyloseq” package (v1.44.0, (44)). Statistically significant differences in alpha diversity between sample groups were assessed using the Kruskal-Wallis test. Beta diversity was visualised by constructing a non-metric multidimensional scaling (nMDS) ordination plot using Bray-Curtis dissimilarity distances. The “vegan” package (v2.6-4, (45)) was used to perform permutational multivariate analysis of variance (PERMANOVA) analysis with 999 permutations on the complete dataset, as well as subsets containing samples from each microbiome fraction to determine significant differences between groups. The core root associated microbiome was defined using the “metagMisc” package (v0.5.0, (46)) and defined as ASVs present in at least 80% of root samples.

### Isolation of *B. distachyon* root associated bacteria

Bacteria isolated from *B. distachyon* roots were initially cultured and maintained on yeast mannitol (YM) agar plates containing (per litre) 2 g mannitol, 3 g D (+) glucose, 0.5 g KH_2_PO_4_, 1 g yeast extract, 0.2 g MgSO_4_·7H_2_O, 0.1 g NaCl, 0.05 g CaSO_4_·2H_2_O and 15 g agar or Luria-Bertani (LB) liquid media containing (per litre): 7.5 g peptone and 2.5 g yeast extract.

Three *B. distachyon* plants were grown in six 7 cm × 15 cm (w × h) pots per soil type, under growth conditions described previously. After 3 weeks, *B. distachyon* plants were harvested and roots were thoroughly washed with sterile 1 × PBS before sonicating for 5 min in an Elma S 300H sonication bath at 37 kHz. Roots were crushed in a sterile mortar and pestle containing 1 mL 1× PBS solution. The resulting solution was serially diluted to 10^-3^ in sterile 1× PBS and used to inoculate growth media. Each dilution was spread onto solid YM agar plates and incubated for 3 days at 25℃. Morphologically distinct colonies were picked and cultured on fresh YM plates at least three times to obtain pure cultures before long-term storage in 25% glycerol at -80℃.

### Screening of isolates for plant growth promoting traits

Only unique, core isolates were screened for plant growth promoting traits (n = 10). To screen isolates for N_2_ fixation, isolates were grown in liquid LB media overnight prior to addition of a loopful of liquid culture to the bottom of 24-well plate wells containing 2 mL of semi solid Nfb media and incubated for 5 d at 28℃. The NFb media contained (per litre): 5 g malic acid, 0.5 g K_2_HPO_4_, 0.2 g MgSO_4_⸱7H_2_O, 0.1 g NaCl, 0.02 g CaCl_2_⸱2H_2_O, 2 mL micronutrient solution (containing (per litre): 0.04 g CuSO_4⸱_5H_2_O, 0.12 g ZnSO_4_⸱7H_2_O, 1.4 g H_3_BO_3_, 1 g Na_2_MoO_4_⸱2H_2_O and 1.175 g MnSO_4_⸱H_2_O), 2 mL bromothymol blue solution (containing (per litre): 5 g bromothymol blue in 0.2N KOH), 4 mL FeEDTA solution (containing (per litre): 16.4 g FeEDTA) and 1 mL vitamin solution (containing (per litre): 0.1 g biotin and 0.2 g pyridoxine HCl). Wells exhibiting the formation of a characteristic bacterial pellicle and a blue colour change indicated nitrogen fixing capacity. Positive wells were used to inoculate fresh wells of semisolid NFb media two additional times to ensure consistency of the nitrogen-fixation capacity of the isolate and reduce instances of false positives. An *Azotobacter* sp. and *Pseudomonas* sp. were used as positive and negative controls, respectively.

To determine the phosphate solubilisation capacity of isolates, National Botanical Research Institute’s phosphate (NBRIP) media containing (per litre): 10 g glucose, 5 g Ca_3_(PO_4_)_2_, 5 g MgCl_2⸱_6H_2_O, 0.25 g MgSO_4_⸱7H_2_O, 0.2 g KCl, 0.1 g (NH_4_)_2_SO_4_ and 15 g agar, was used. To determine the capacity of isolates to produce siderophores, chromeazurol-S (CAS) media was used. The CAS media consisted of an LB agar base (7.5 g tryptone, 2.5 g yeast extract, 15g agar per litre) and CAS solution (1.21 g chromeazurol-S, 1.82 g CTAB and 0.27 g FeCl_3_⸱6H_2_O per litre), mixed in a 9:1 (LB:CAS) ratio. Cultures were diluted using liquid LB to reach a final optical density at 600 nm (OD600) of 0.1. Five 10 µL droplets were added to a NBRIP and CAS plate and incubated at 25℃ for 7 and 3 days, respectively. Isolates were considered positive for siderophore production if a zone of colour change from blue to orange was observed around colonies. Isolates of interest were grown overnight in liquid LB media. Isolates were considered positive for phosphate solubilisation upon observing a zone of clearing around colonies. A phosphate solubilisation index (PSI) was calculated by dividing the diameter of a zone of clearing by the diameter of the bacterial colony.

For screening of indole acetic acid production, isolates were grown in liquid TSB media amended with 2.5 mM tryptophan (17 g tryptone, 3 g peptone, 5 g NaCl, 2.5 g KH_2_PO_4_, 2.5 g dextrose and 0.5 g L-tryptophan per litre) and quantified through the use of Salkowski’s reagent consisting of 2 mL of a 0.5 M FeCl_3_⸱6H_2_O solution, 49 mL dH_2_O and 49 mL 35% (v/v) HClO_4_. A pre–inoculum of isolates of interest were grown overnight in liquid TSB media. Cultures were then standardized to an OD_600_ of 0.1 and used to inoculate fresh liquid TSB media. These cultures were incubated overnight in the dark before centrifuging and mixing supernatant in a 1:1 ratio with Salkowski’s reagent. Presence of indole content in samples was determined by measuring the colour change at 530 nm using an xMark^TM^ microplate absorbance spectrophotometer (Bio-Rad).

### Identification of bacterial isolates

To identify isolates, their 16S rRNA gene was amplified using 27F (5′-AGAGTTTGATCMTGGCTCAG′) and 1429R (5′-TACGGYTACCTTGTTACGACTT′) primers (47). PCR consisted of 32 cycles of 98℃ for 10 s, 62℃ for 30 s and 72℃ for 60 s before a final extension at 72℃ for 5 min. Samples were sent to AGRF in Melbourne, Victoria for Sanger sequencing. Isolates were identified using the Basic Local Alignment Search Tool (BLAST) to compare 16S rRNA gene sequences against the NCBI nucleotide database.

### Plant growth promotion assay

A gnotobiotic, semi hydroponic growth system was adapted and used to assess the plant growth promotion capacity of selected bacterial isolates (48). Magenta™ box model GA-7 was used as the system housing. Boxes were filled to 2 cm with 10% HCl washed and rinsed, autoclaved polypropylene pellets (LyondellBasell) to hold plants in place. Pellets were saturated with 20 mL of ¼ strength Hoagland nutrient solution containing macronutrients as follows: 5 mM Ca(NO_3_)_2_, 5 mM KNO_3_, 2 mM MgSO_4_, 41.5 µM FeEDTA(Na), 1 mM KCl, 1 mM KH_2_PO_4_ and micronutrients as: 92 µM H_3_BO_3_, 20 µM MnCl_2_, 1.6 µM ZnCl_2_, 0.5 µM CuCl_2_ and 0.24 µM Na_2_MoO_4_ (49). Hoagland’s solution was buffered to pH 6.5 with 1 mM MES-hydrate buffer.

Dehusked and surface sterilised *B. distachyon* seeds were imbibed for 2 d at 4°C in the dark prior to placing on 0.9% agar plates for germination at room temperature for 3 days in the dark. Three germinated seeds were placed on top of polypropylene pellets in each box, slightly beneath the surface of the nutrient solution to ensure roots did not dry out. Each box represented one biological replicate. Control and treatments were replicated four times. Plants were grown for 4 d in a 18/6 day/night cycle at 24℃ and 18℃, respectively, in a Conviron growth cabinet. After four days, plants were inoculated with selected isolates that were grown overnight in LB liquid media. To ensure there was no residual growth media transfer, isolates were pelleted and washed with sterile Hoagland’s solution, prior to adjustment to an OD_600_ of 0.2. Inoculant (50 µL) was added to the base of each seedling shoot. Twenty-one days post inoculation (DPI), plants were harvested and oven dried at 70℃ for 48 h to obtain shoot and root dry weights. Statistical differences between group means were calculated using an unpaired t-test in in R (v4.4.0) and R Studio (v2023.06.0) (43).

## Data availability statement

All raw amplicon sequencing reads for microbial community analysis are available on the NCBI Sequence Read Archive under accession number PRJNA1120948.

## Acknowledgements

This work was supported by the Australian Research Council through the Industrial Transformation Research Hub Scheme (grant number IH200100023) awarded to H-W. Hu, U. Roessner and D. Chen.

## Author Contribution Statement

C.O.P, Z.F.I and H-W.H conceived the study. C.O.P and Z.F.I collected soils samples. C.O.P conducted experiments. E.C assisted with laboratory work. C.O.P and L.H.C performed data analysis. C.O.P, H.L.H, E.C, Z.F.I and H-W.H provided valuable insight and discussion. C.O.P, Z.F.I and H-W.H wrote the manuscript. All authors provided critical feedback and final approval prior to publication.

## Appendix

**Figure A1:**
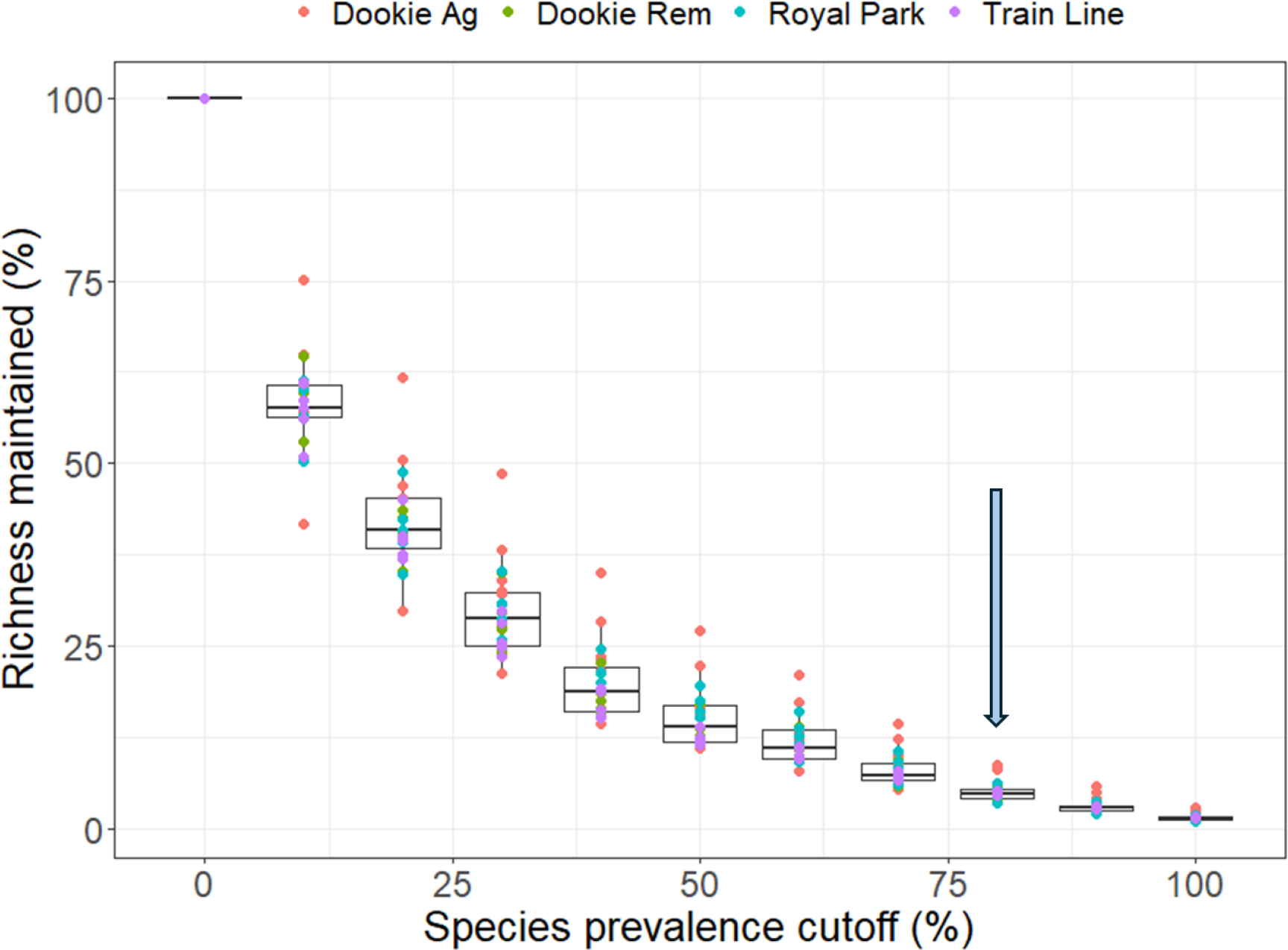
Richness maintained in root fractions at species prevalence cutoffs at 10% intervals. An 80% prevalence threshold was selected for definition of the core *Brachypodium distachyon* root-associated microbiome (blue arrow).

**Figure A2:**
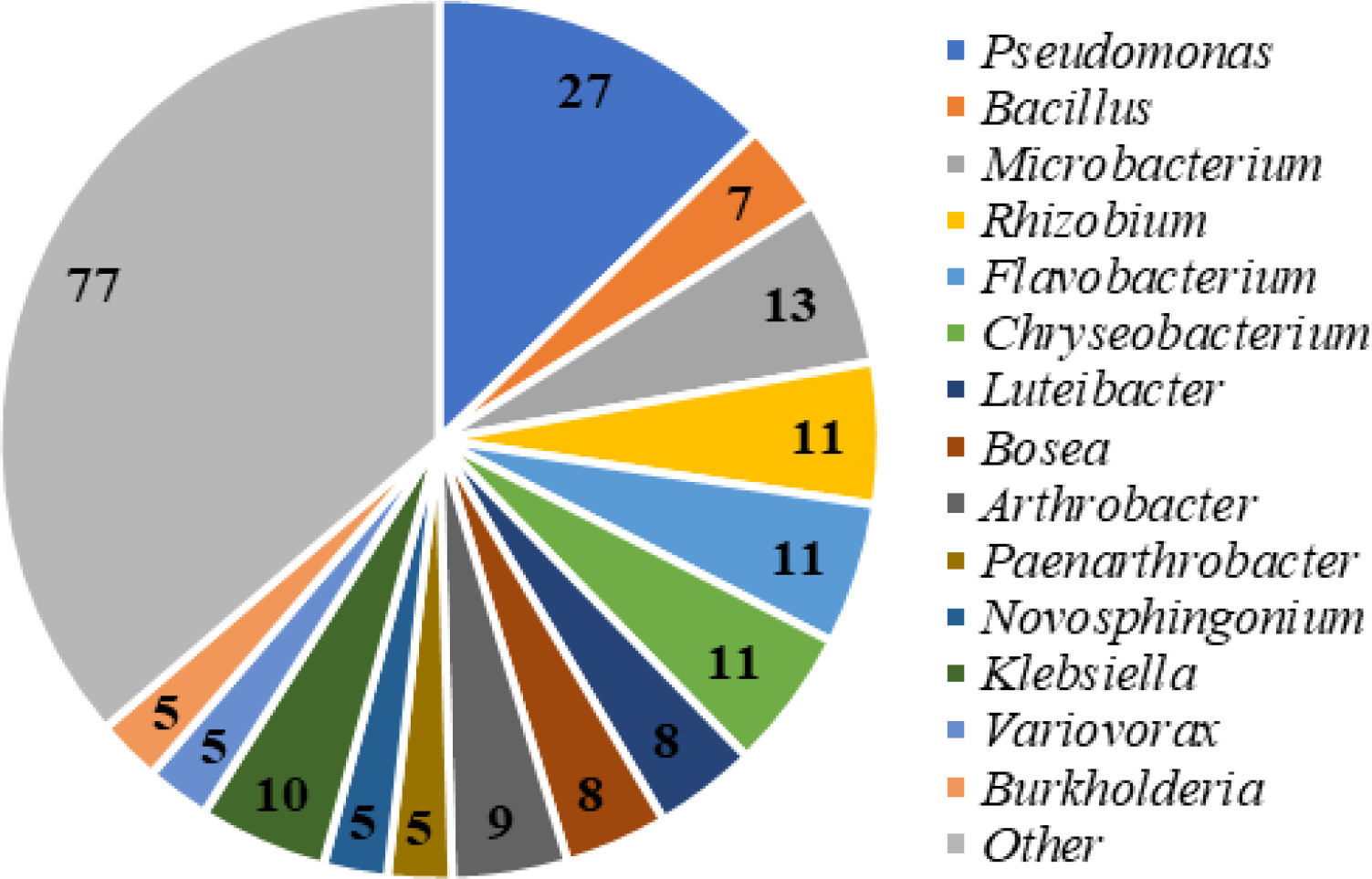
Frequency of bacterial genera isolated from *B. distachyon* roots.

**Table A1:**
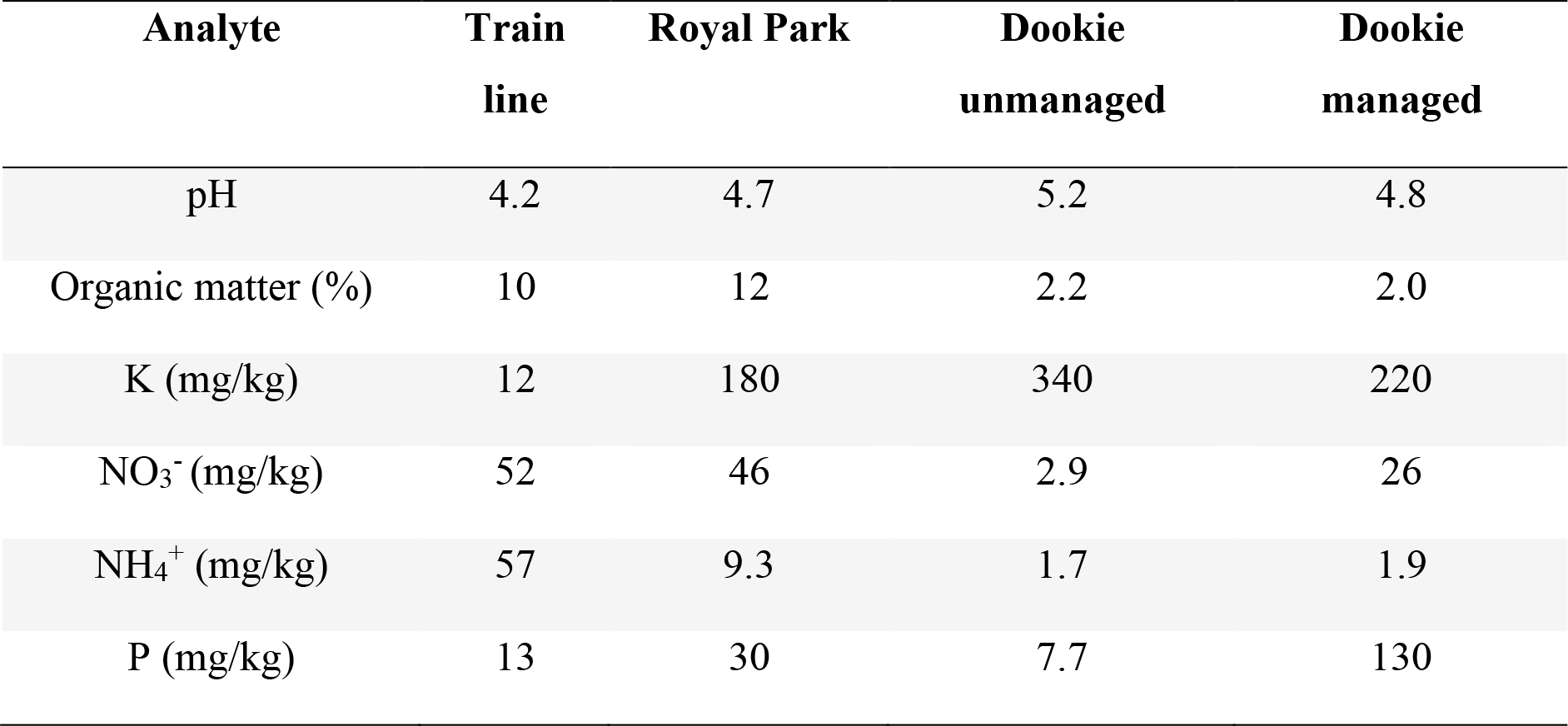
Key physicochemical properties of collected soils.

**Table A2:** Factors contributing to microbiome diversity determined via PERMANOVA analysis using the Bray-Curtis metric with 999 permutations.

|  | **All** |  | **Bulk** |  | **Rhizosphere** |  | **Root** |  |
| --- | --- | --- | --- | --- | --- | --- | --- | --- |
| **Factor** | *F*- value | R<sup>2</sup> | *F*- value | R<sup>2</sup> | *F*- value | R<sup>2</sup> | *F*- value | R<sup>2</sup> |
| Location | 14.68 | 39.7* | 21.2 | 76.0* | 17.9 | 72.8* | 7.2 | 53.2* |
| Fraction | 6.6 | 16.3* | - | - | - | - | - | - |

**Table A3:** Mean root:shoot ratio, leaf number and tiller number of *B. distachyon* inoculated with core *B. distachyon* root associated isolates (n = 4).

| Isolate (BD#) | Origin<sup>a</sup> | Root:Shoot | Leaf number |
| --- | --- | --- | --- |
| Control | N/A | 0.31 ± 0.06 | 6.75 ± 1.07 |
| *Neobacillus* BD17 | TL | 0.28 ± 0.10 | 6.42 ± 1.50 |
| *Bacillus* BD33 | TL | 0.43 ± 0.08 | 8.67 ± 0.61* |
| *Bacillus* BD87 | RP | 0.39 ± 0.07 | 7.67 ± 0.23 |
| *Bacillus* BD101 | RP | 0.38 ± 0.06 | 7.91 ± 0.99 |
| *Bacillus* BD115B | RP | 0.40 ± 0.06 | 6.42 ± 1.50 |
| *Bosea* BD19 | TL | 0.47 ± 0.04* | 6.67 ± 1.98 |
| *Paraburkholderia*<br>BD210 | DU | 0.50 ± 0.21 | 8.25 ± 0.71 |
| *Burkholderia*<br>BD284 | DM | 0.38 ± 0.03 | 7.08 ± 1.31 |
| *Leifsonia* BD292 | DM | 0.39 ± 0.08 | 7.75 ± 1.31 |
| *Microbacterium*<br>BD86 | RP | 0.36 ± 0.05 | 6.41 ± 1.30 |
| *Rhodococcus*<br>BD157 | DU | 0.40 ± 0.12 | 6.42 ± 1.50 |
| *Rhodococcus*<br>BD285 | DM | 0.42 ± 0.07 | 8.25 ± 0.71 |

<sup>a</sup> RP = Royal Park, TL = Train line, DM = Dookie managed, DU = Dookie unmanaged
\* = p < 0.05

